# LRRK2 is recruited to phagosomes and co-recruits Rab8 and Rab10 in human pluripotent stem cell-derived macrophages

**DOI:** 10.1101/779835

**Authors:** Heyne Lee, Rowan Flynn, Ishta Sharma, Phillippa J. Carling, Francesca J. Nicholls, Monika Stegmann, Jane Vowles, Richard Wade-Martins, William S. James, Sally A. Cowley

## Abstract

The Parkinson’s disease-associated gene, *LRRK2*, is also associated with immune disorders and infectious disease, and is expressed in immune subsets. Here, we characterise a platform for interrogating the expression and function of endogenous LRRK2 in authentic human phagocytes, using human induced Pluripotent Stem Cell-derived macrophages and microglia. Endogenous LRRK2 is expressed and upregulated by interferon-γ in these cells, including a 187kD cleavage product. Using LRRK2 knockout and G2019S isogenic repair lines, we find that LRRK2 is not involved in initial phagocytic uptake of bioparticles, but is recruited to LAMP1(+)/Rab9(+) ‘maturing’ phagosomes, and LRRK2 kinase inhibition enhances its residency at the phagosome. Importantly, LRRK2 is required for Rab8a and Rab10 recruitment to phagosomes, implying that LRRK2 operates at the intersection between phagosome maturation and recycling pathways in these professional phagocytes.

## Introduction

*LRRK2* (leucine-rich repeat kinase 2) encodes a large (286 kD), multi-domain cytoplasmic protein, with both GTPase and kinase domains, flanked by several protein-protein interaction domains. Mutations in *LRRK2* account for approximately 4% of familial and 1% of idiopathic cases of the progressive neurodegenerative disorder, Parkinson’s disease, forming an important genetic risk factor for PD. Most PD-causing mutations cluster within the two enzymatic sites, notably G2019S and R1441C/G (leading to a modest 2-3 fold increased kinase activity, and decreased GTPase activity, respectively) (Ferreira and Massano, 2017). *LRRK2* variants are also associated with autoimmune disorders(Witoelar et al., 2017), particularly Crohn’s Disease, and with infectious diseases, notably *Mycobacterium leprae*(Wang et al., 2015; Zhang et al., 2009). *LRRK2* expression has also been linked with *Mycobacterium tuberculosis* infection(Härtlova et al., 2018; Wang et al., 2018). *LRRK2* is expressed in a variety of cell lineages, including several immune subsets, notably B cells, neutrophils, monocytes, macrophages and microglia(Atashrazm et al., 2018; Fan et al., 2018; Gardet et al., 2010; Hakimi et al., 2011; Kim et al., 2012; Marker et al., 2012; Moehle et al., 2012; Thévenet et al., 2011) (reviewed in (Lee et al., 2017)).

Macrophages populate most tissues of the body, deriving initially from primitive macrophages that migrate in during embryogenesis, and are replenished as necessary during the lifespan by either local proliferation and/or replacement by blood monocyte-derived macrophages, depending on the tissue (Hoeffel and Ginhoux, 2015). Macrophages perform tissue homeostatic functions and are also a first-line defence against pathogens, armed with a plethora of pattern-recognition and opsonin receptors. They rapidly phagocytose and kill incoming bacteria, fungi and protoctists, and have strong antiviral defences. Nonetheless, various pathogens can overcome these defences to survive and proliferate in macrophages, including *M.leprae* and *M.tuberculosis*, Interestingly, LRRK2 has recently been shown to be required for survival of *M.tuberculosis* in macrophages(Härtlova et al., 2018).

Microglia are a resident, primitive macrophage-derived population in the central nervous system (CNS), performing homoeostatic functions (phagocytosing cell debris, extracellular protein aggregates and incompetent synapses) to maintain a healthy environment for neurons. However, they can also secrete inflammatory mediators when activated, notably TNFα, and a myriad of cytotoxic factors, especially reactive oxygen species (ROS) and nitric oxide (NO), which can instigate a feedforward cycle of chronic inflammation and neurodegeneration. Therefore, microglia are not only involved in preventing neurodegenerative disease by phagocytosing potentially harmful materials but also can contribute to disease progression by initiating exaggerated inflammatory responses (reviewed in (Wolf et al., 2017)).

Due to the difficulty in obtaining primary patient material, most studies of LRRK2 have used either animal models, *in vitro* biochemical assays, or transformed cell lines, often involving non-physiological exogenous overexpression of LRRK2 in irrelevant lineages. Such observations need to be subsequently validated in a karyotypically normal human cellular system at physiologically relevant expression levels to demonstrate their applicability to normal human physiology and disease. We have previously developed methods for efficient differentiation of human induced pluripotent stem cells (hiPSC) to macrophages, which exhibit authentic phagocytic properties and cytokine-profiles(Flynn et al., 2015; Haenseler et al., 2017b; Karlsson et al., 2008; van Wilgenburg et al., 2013). The differentiation pathway is demonstrably independent of c-Myb expression(Buchrieser et al., 2017), indicating that they represent an embryonic/primitive ontogeny, and are therefore also suitable as a precursor for differentiation to microglia. We have shown that they can be further differentiated to microglia by co-culture with hiPSC-neurons, whereupon they acquire a ramified morphology and associated neuronal surveillance activity(Haenseler et al., 2017a).

In this study we have used hiPSC-macrophages and microglia from patient, control and gene-edited lines to explore expression of LRRK2 protein from the endogenous locus and the role of LRRK2 in this lineage. We show that LRRK2 is expressed in hiPSC-macrophages and microglia, with expression significantly upregulated by interferon-γ (IFNγ), and identify the cleavage region of a truncated LRRK2 product found in this lineage. In this system, LRRK2 is not involved in the initial phagocytic uptake of particles, but is recruited to maturing phagosomes, and this is exacerbated by inhibition of LRRK2 kinase activity. Importantly, we show that LRRK2 is required for recruitment to phagosomes of Rab8a and Rab10 (members of the membrane trafficking regulator family of Rab GTPases and substrates of LRRK2 kinase activity). This demonstrates that LRRK2 operates at the intersection between phagosome maturation and recycling pathways in the myeloid lineage.

## Results

### Characterisation of LRRK2 Knockout and G2019S Isogenic control hiPSC lines and a major LRRK2 cleavage product in macrophages

The hiPSC lines used in this study are listed in Table S1, with quality control information in Figure S1. WT.1 – WT.6 were from six healthy control donors.

*LRRK2* knockout was generated in a control hiPSC (line WT.1) by a double nickase CRISPR/Cas9(Ran et al., 2013) strategy, using a pair of gRNAs targeting exon 3 of *LRRK2* (Figure 1A and S2A). A patient line containing a heterozygous *LRRK2* mutation G2019S (GS) was successfully repaired to WT (GS-Repair) as shown by sequence analysis (Figure 1B). Two KO clones (KO.1 and KO.2) displayed out-of-frame homozygous deletion of *LRRK2* (Figure S1A) and showed complete absence of LRRK2 protein when differentiated to macrophages (Figure 1C). There was no significant difference in the production of macrophage precursors in edited versus parental lines (Figure S2B and C).

**Figure 1.**
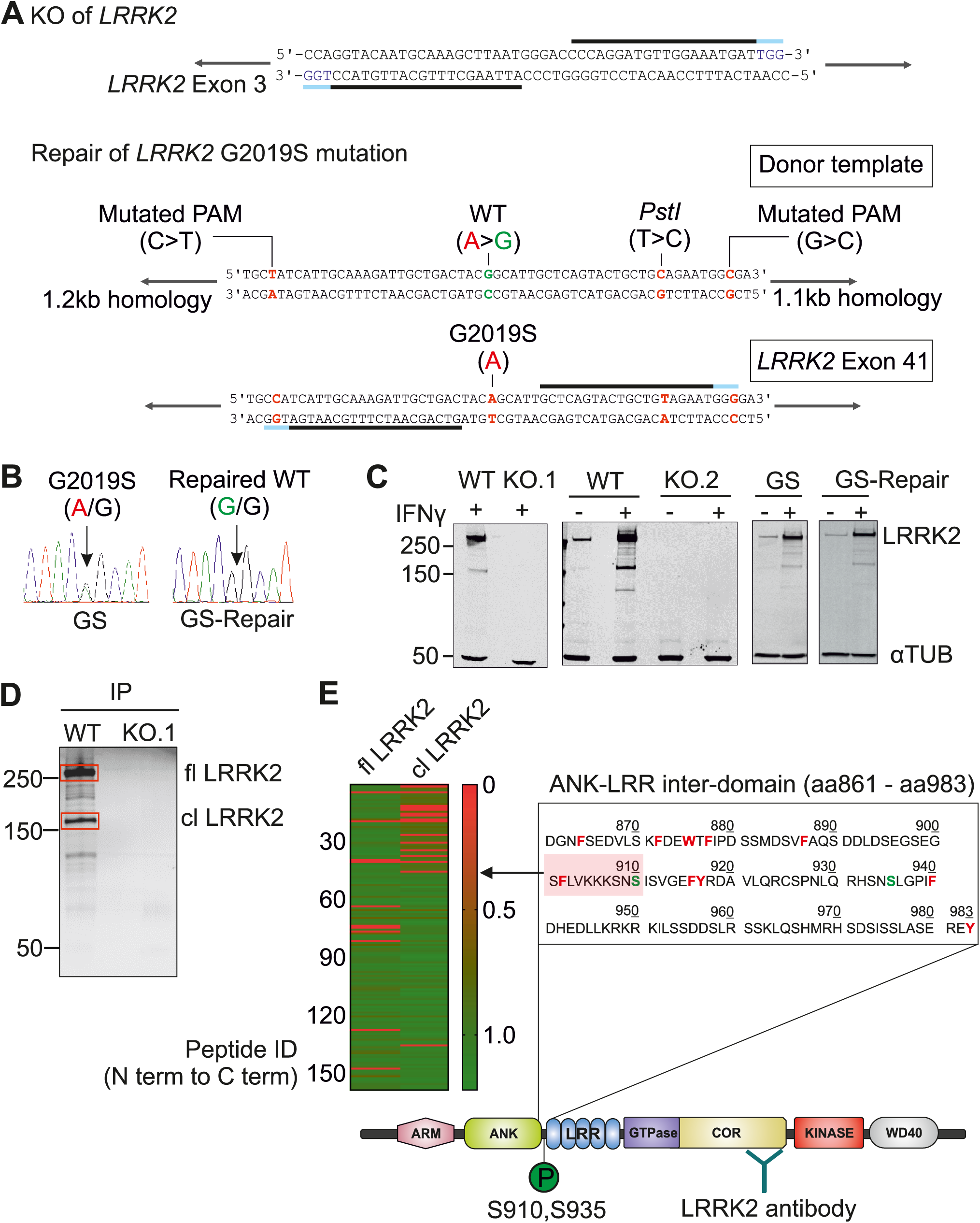
Characterisation of LRRK2 Knockout and G2019S Isogenic control hiPSC lines and a major LRRK2 cleavage product in macrophages. (A) CRISPR/Cas-9 mediated knockout (KO) of *LRRK2* was performed using a double nickase strategy with a pair of guide RNAs (gRNAs, black lines, PAM, blue) targeting exon 3 of *LRRK2*. For CRISPR/Cas-9-mediated repair of *LRRK2* G2019S mutation, the donor template contained silent mutations in the PAM site to maximize gene-editing efficiency and a *PstI* site for the subsequent screening of edited clones. (B) Sequencing results demonstrate that the G2019S mutation present in one allele (A/G) was successfully corrected to the WT sequence (G/G). (C) Western blot probed with antibody to LRRK2 N241A/34 shows the complete absence of endogenous LRRK2 protein in both KO clones in hiPSC-macrophages with or without IFNγ (100 ng/mL) activation. The total expression level of LRRK2 protein in the isogenic repaired line is not altered. (D) Western blot showing immunoprecipitated (N241A/34 antibody) endogenous LRRK2 protein from WT and KO derived hiPSC-macrophages. Silver-stained gel bands corresponding to full-length LRRK2 (fl LRRK2) and the cleaved LRRK2 (cl LRRK2), indicated in red boxes, were cut and analyzed by mass spectrometry. (E) Trypsin-digested peptides were quantified by MaxQuant. Heat map shows the intensity of identified peptides (ordered from N-terminal to C-terminal) of fl LRRK2 and cl LRRK2. cl LRRK2 is largely missing N-terminal peptides up to the region aa 901 – 910, indicated by pink box in schematic (Chymotrypsin cleavage sites, red; phosphorylation sites, green).

Western blot of hiPSC-macrophages using LRRK2 monoclonal antibody N241A/34 (binding site amino acids 1836-1845) consistently showed multiple faint bands and a major band (approximately 170 kD relative to size markers) in addition to full-length LRRK2 (286 kD). This was particularly evident upon interferon-γ stimulation. The staining pattern was confirmed to be *LRRK2*-specific by its absence in LRRK2 KO hiPSC-macrophages (Figure 1C). Since the 170 kD band was not evident when probed with an N-terminal antibody (N138/6) (Figure S2D) and LRRK2 transcripts have not been reported that would correspond to this product, we reasoned that this is likely a proteolytically cleaved product. It is not a result of technical post-lysis proteolysis, as spiking recombinant full length LRRK2 into *LRRK2*-KO macrophage lysate did not lead to its proteolytic degradation under our standard lysis conditions (Figure S2D). Addition of protease inhibitors to live macrophages reduced the proportion of the truncated versus full-length protein, implying that it is a natural cleavage generated within intact macrophages (Figure S2E). Immunoprecipitation using the N-terminal antibody revealed that the cleaved product can heterodimerize with full-length LRRK2 (Figure S2F). To identify the cleavage site, we isolated endogenous LRRK2 protein from hiPS-macrophages by immunoprecipitation with N241A/34 antibody, ran it on a denaturing gel and analysed the cleaved product by mass spectrometry (Figure 1D). MaxQuant analysis of trypsin-digested peptide fragments revealed that cleavage occurs within the ANK-LRR interdomain region (aa861 – aa983), particularly between aa901 – aa910, (which contains a predicted chymotrypsin cleavage site), generating a C-terminal product of 1626aa (predicted 185 kD) (Figure 1E).

### IFNγ increases LRRK2 protein expression in hiPSC-macrophages and microglia

IFNγ has been shown to upregulate LRRK2 protein expression in myeloid cells(Gardet et al., 2010). Similarly, in hiPSC-macrophages, LRRK2 protein expression increased significantly (up to 10-fold) upon IFNγ treatment (Figure 2A and Figure S3A). Phosphorylation of LRRK2 at S935 was observed, significantly decreasing in the presence of LRRK2 kinase inhibitor GNE-7915, in accordance with the published literature(Hatcher et al., 2017) (Figure 2B). hiPSC-macrophages with the heterozygous G2019S mutation showed the same pattern, with no significant difference in either the basal phosphorylation level at S935 or the degree of dephosphorylation upon treatment with LRRK2 kinase inhibitors compared to its isogenic pair (Figure S3B), likely because G2019S only produces a modest 2-fold increase in kinase activity.

**Figure 2.**
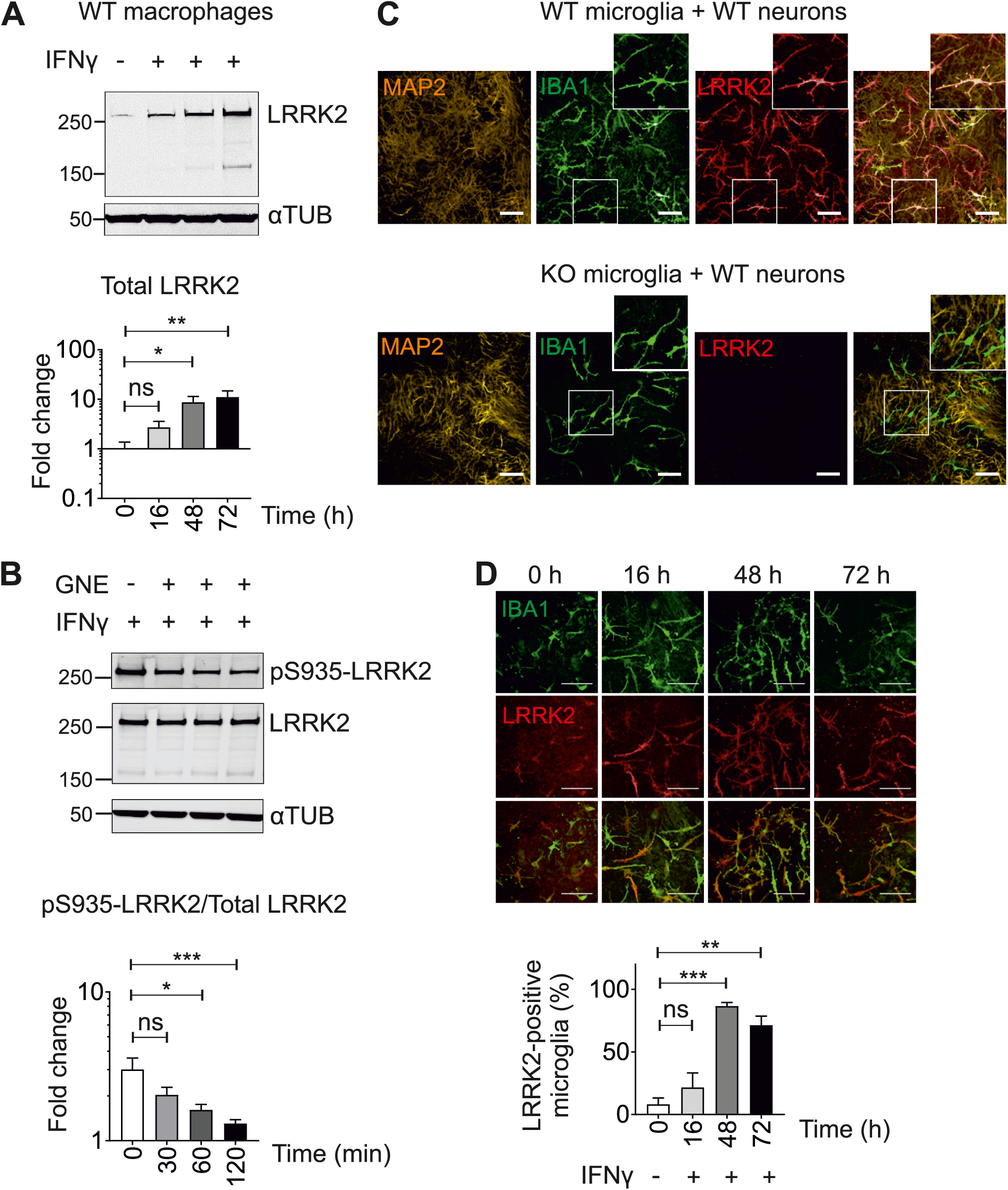
IFNγ increases LRRK2 protein expression in hiPSC-macrophages and microglia. (A) hiPSC-macrophages were treated with IFNγ (100 ng/mL) for 16, 48, and 72 h and cell lysates analyzed by western blot. Bar graph shows fold change of the total endogenous LRRK2 protein over loading control (αTUB; alpha-tubulin). (B) hiPSC-macrophages were treated with LRRK2 kinase inhibitor GNE-7915 (GNE; 1 µM) for 30, 60, and 120 min after 72 h treatment with IFNγ. Bar graph shows fold change in pS935-LRRK2 signal over total LRRK2 relative to DMSO control. All data points represent mean ± SEM of at least three independent experiments. (C) Microglia differentiated from either WT or KO hiPSC were co-cultured with cortical neurons differentiated from WT hiPSC. Cells were treated with IFNγ (100 ng/mL) for 48 h and were stained with antibodies against neuronal marker (MAP2), microglial marker (IBA1) and LRRK2 (N241A/34). Scale bars represent 100 µm. (D) hiPSC-microglia co-cultured in hiPSC-neurons were treated with IFNγ (100 ng/mL) for 0, 16, 48, and 72 h. Z-stacked confocal images were acquired and the number of hiPSC-microglia (IBA1-positive cells) expressing LRRK2 was quantified using Columbus software (PerkinElmer). Bar graph shows percent LRRK2-expressing hiPSC-microglia as mean ± SEM of three independent experiments. Statistical significance was tested using one-way ANOVA.

We next examined LRRK2 expression in hiPSC-microglia, co-cultured with hiPSC-cortical neurons. LRRK2 protein was clearly expressed in hiPSC-microglia, while its expression level was not detectable in hiPSC-cortical neurons (Figure 2C). The specificity of LRRK2 staining in microglia was confirmed by co-culturing microglia differentiated from LRRK2 KO hiPSC with cortical neurons differentiated from LRRK2 WT hiPSCs (Figure 2C). To test whether IFNγ upregulates LRRK2 protein in hiPSC-microglia or neurons, hiPSC-microglia/cortical neuron co-cultures were treated with IFNγ for 16 h, 48 h, or 72 h. IFNγ treatment significantly upregulated the percent of LRRK2 expressing hiPSC-microglia to 86%, its expression level plateauing by 48 h post IFNγ treatment (Figure 2D).

Together, these results demonstrate the validity of the hiPSC-macrophage and microglia models for investigating endogenous LRRK2 function.

### LRRK2 is not involved in the initial phagocytic uptake of bioparticles but is recruited to maturing phagosomes

We next investigated whether LRRK2 is involved in phagocytosis using hiPSC-macrophages. hiPSC-macrophages readily phagocytose a wide variety of ‘meals’, including killed yeast bioparticles (‘zymosan’). Complete absence of LRRK2 in hiPSC-macrophages did not alter their ability to take up fluorescent zymosan, with or without IFNγ induction (Figure 3A, B). Similarly, zymosan uptake by G2019S patient derived hiPSC-macrophages was not significantly different from that of its isogenic pair (Figure 3C). Lastly, pharmacological inhibition of LRRK2 kinase activity with two structurally distinct LRRK2 kinase inhibitors, GSK2578215A (GSK) or GNE-7915 (GNE), had no significant effect on zymosan uptake (Figure 3D). Acidification of the phagosomes, as assessed by uptake of pH-sensitive fluorescent (pHrodo) zymosan particles, was also not significantly altered by manipulating LRRK2 in hiPSC-macrophages (Figure S4).

**Figure 3.**
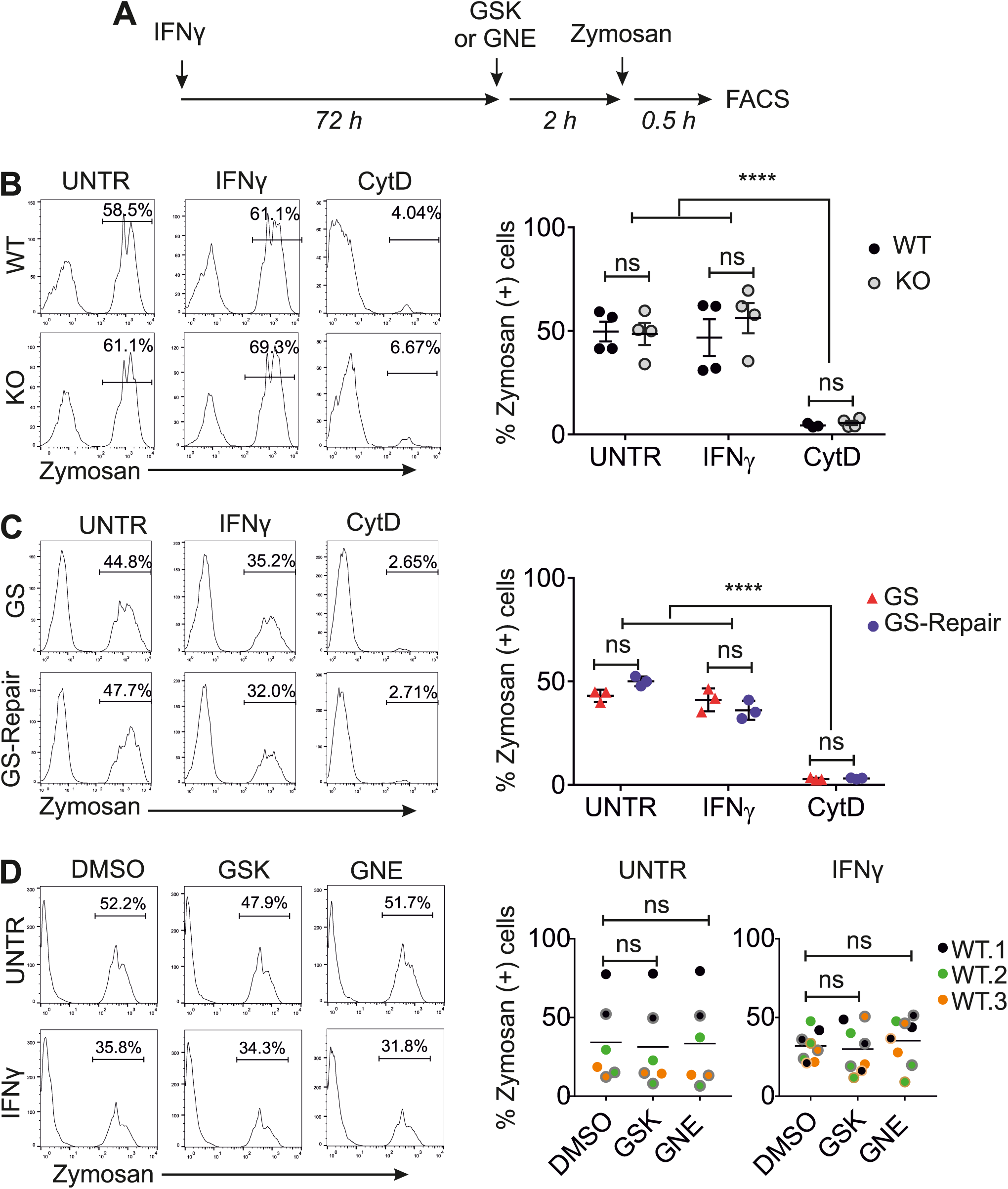
LRRK2 is not involved in the initial phagocytic uptake of bioparticles. (A) Outline of the experimental design. (B and C) hiPSC-macrophages differentiated from the two isogenic pairs (B, WT and KO; C, GS and GS-Repair) were treated with IFNγ for three days, then pre-treated with either DMSO or Cytochalasin D (CytoD; 10 µM) for 1 h, then incubated with Alexa-488 conjugated zymosan bioparticles for 30 min. Bar graphs show the percentage of zymosan(+) cells as mean ± SEM. (D) hiPSC-macrophages from three healthy controls (WT.1, WT.2, WT.3) were pre-treated with LRRK2 kinase inhibitors (GNE or GSK; 1 µM) prior to the addition of zymosan bioparticles. Each dot represents data from each independent experiment. Paired t-test was used for statistical analysis.

While no functional difference in initial phagocytic uptake was observed across LRRK2 lines, confocal imaging clearly demonstrated the presence of LRRK2 on a subset of zymosan-containing phagosomes (in IFNγ treated cells to enable visualisation of LRRK2) (Figure 4A). This was also observable with *E.coli* bioparticles, with and without opsonization, and *S.typhimurium*, (Figure 4A). LRRK2 was not observed on phagosomes containing αsyn fibrils, even when opsonised (Figure S5), indicating that LRRK2 recruitment is context-dependent. The number of LRRK2 positive zymosan-containing phagosomes was time-dependent, peaking at 1-2 h post-addition of the meal to cells (mean 14.7%, range 6.8-25.2% at 2 h), so 2 h zymosan incubation was used for all subsequent experiments (Figure 4B). LRRK2-positive (LRRK2+) phagosomes were found to also be positive for the late phagosomal markers lysosome-associated protein LAMP-1 and Rab9, with significantly more LRRK2(+)LAMP-1(+) or LRRK2(+)Rab9(+) phagosomes than LRRK2(+)Rab5(+) phagosomes (an early phagosome marker) (Figure 4C). Together, this shows that LRRK2 is recruited during later stages of phagosome maturation in hiPSC-macrophages, around the time when lysosomes are recruited to phagosomes.

**Figure 4.**
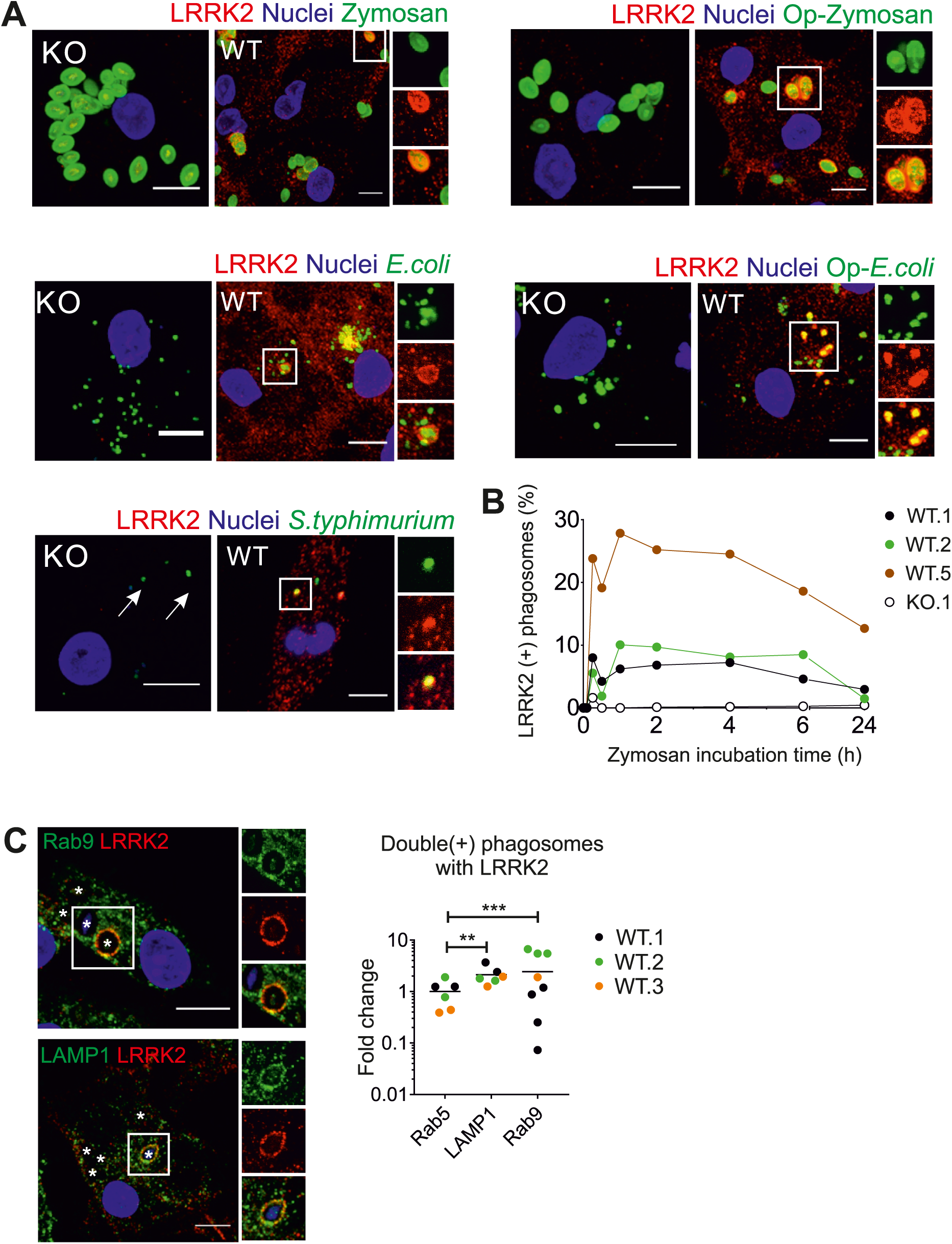
LRRK2 is recruited to maturing phagosomes. (A) Confocal images of LRRK2 WT and LRRK2 KO hiPSC-macrophages fed with zymosan, IgG-opsonized zymosan (Op-Zymosan), *E.coli*, IgG-opsonized *E. coli* (Op-*E. coli)*, GFP-expressing *S. typhimurium*, (all Alexa-488-conjugated). Phagocytosis proceeded for 2 h before cells were fixed and stained with antibody against LRRK2. All scale bars represent 10 µm. (B) Quantification of LRRK2(+) phagosomes over time. Each data point represents the percentage of total phagosomes that were LRRK2 positive (by automated image analysis), mean of 20 random fields. Note, 15 minute timepoint accuracy is poor due to low absolute numbers of phagosomes at this early timepoint. (C) hiPSC-macrophages fed with zymosan bioparticles were stained with antibodies against LRRK2 and markers associated with maturing phagosomes, LAMP1 and Rab9. Internalized zymosan bioparticles are indicated by an asterix to better visualize decoration of phagosome membrane by LRRK2 and marker staining. Bar graph shows fold change in the number of LRRK2(+) phagosomes displaying late phagosomal markers versus early phagosome marker (Rab5). Student t-test was used for statistical analysis.

### Rab8a and Rab10 are recruited to phagosomes and this is LRRK2-dependent

Rab GTPases regulate various fission and fusion events during phagocytosis, and in LRRK2-overexpression systems and non-human systems LRRK2 has been shown to associate physically with subsets of this family of proteins (Dodson et al., 2011; Gómez-Suaga et al., 2014; Steger et al., 2016; Waschbüsch et al., 2014; Yun et al., 2015). Importantly, several Rabs, particularly Rab8a and Rab10, have been identified as physiological substrates of LRRK2, able to be phosphorylated by LRRK2 (on Thr72/73 respectively)(Fan et al., 2018; Steger et al., 2016). We therefore investigated whether these Rab GTPases are involved during LRRK2 recruitment to phagosomes. Rab8a and Rab10 could be observed coating the same phagosomes as LRRK2, while Rab7 had no significant association with LRRK2(+) phagosomes (Figure 5A,B). 14-3-3 proteins, which associate with pS935-LRRK2(Dzamko et al., 2010), did not appear closely colocalised with LRRK2, but could be observed just peripheral to LRRK2-coated clusters of phagosomes (not quantified). Importantly, the number of Rab8(+) and Rab10(+) phagosomes was significantly reduced (to background levels) in LRRK2 KO hiPSC-macrophages (Figure 5A,C), demonstrating that the presence of LRRK2 is necessary for recruitment of Rab8a and Rab10 to phagosomes.

**Figure 5.**
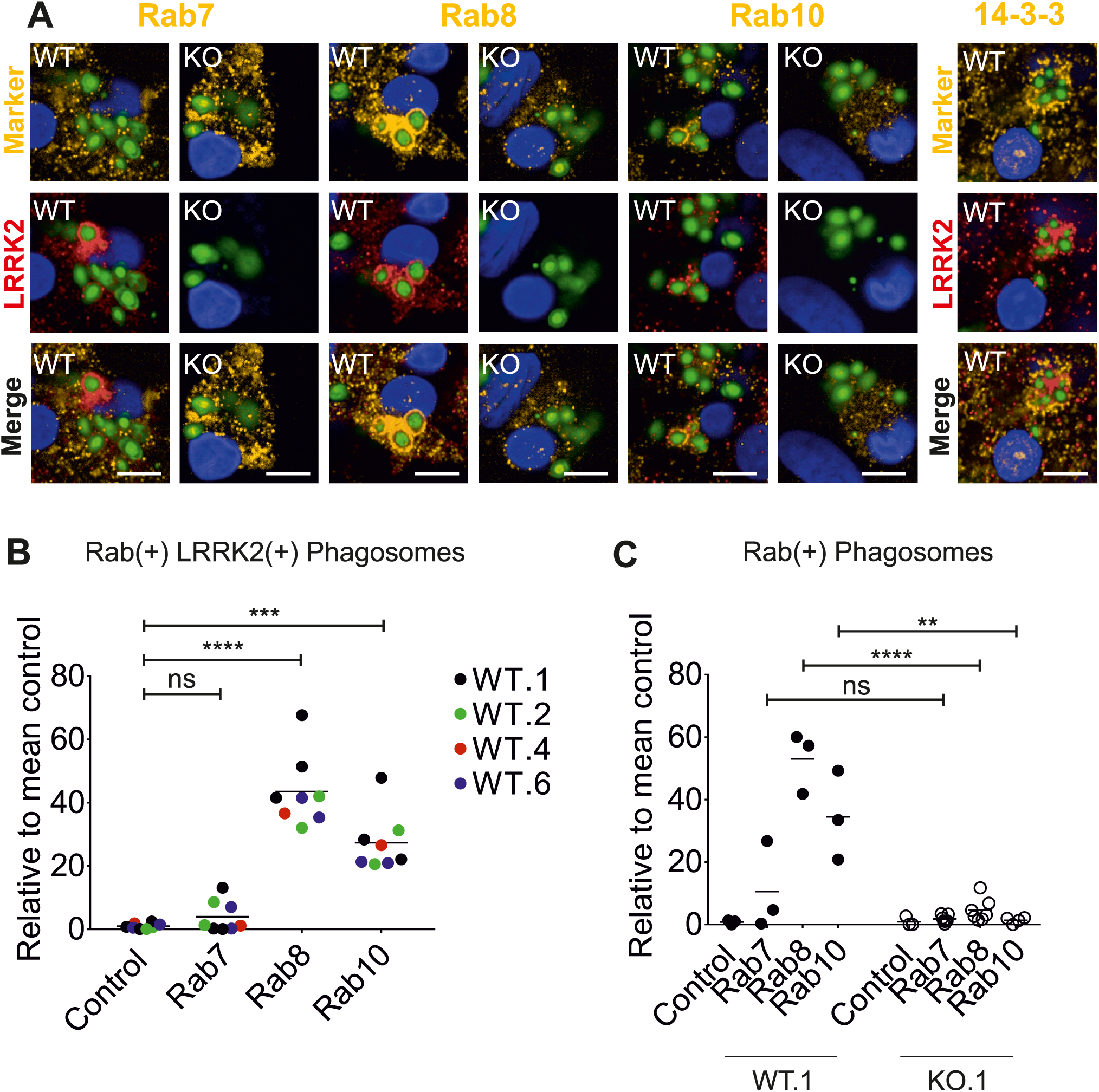
Rab8 and Rab10 are recruited to phagosomes and this is LRRK2-dependent. (A) Confocal images of LRRK2 WT and LRRK2 KO hiPSC-macrophages fed with zymosan Alexa-488 for 2 h, then fixed and stained for LRRK2 (Alexa-647, red) and Rabs or pan-14-3-3 protein (Alexa-546, yellow). Scale bars represent 10 µm. (B) Quantification of number of LRRK2(+) phagosomes that are also Rab(+). Data from four WT hiPSC-macrophage lines (WT.1, WT.2, WT.4, WT.6), each data point represents the mean of 20 fields per independent replicate. Control, secondary antibody only. 1-way ANOVA with Tukey’s multiple comparison was performed for statistical analysis. (C) Quantification of Rab(+) phagosomes, each data point represents the mean of 20 fields per independent replicate. Significance assessed by Student t-test.

### LRRK2 kinase inhibitors increase LRRK2 density at phagosomes, but inhibit Rab8a and Rab10 recruitment

In LRRK2-overexpressing HEK 293T cells, it has been reported that LRRK2 kinase inhibitors affect localization of LRRK2 within the cell, from diffused cytosolic distribution to more discrete cytosolic pools(Dzamko et al., 2010). Therefore, we investigated whether the application of LRRK2 kinase inhibitors would affect LRRK2 recruitment to phagosomes during phagocytosis. Pre-treating hiPSC-macrophages with LRRK2 kinase inhibitors did not change the total number of LRRK2(+) phagosomes. However, we noticed that significantly more (four fold) of these LRRK2(+) phagosomes displayed enhanced LRRK2 signal (referred as ‘super-coated’ LRRK2 phagosomes, LRRK2(++), Figure 6A-C). This was also observable with the LRRK2 G2019S patient line and its isogenic control (Figure S6A,B). There was no significant difference between the isogenic pair of lines, suggesting that monoallelic G2019S LRRK2 kinase enhancement is not potent enough to give a detectable difference in ‘super-coating’ compared with strong drug-induced kinase inhibition.

**Figure 6.**
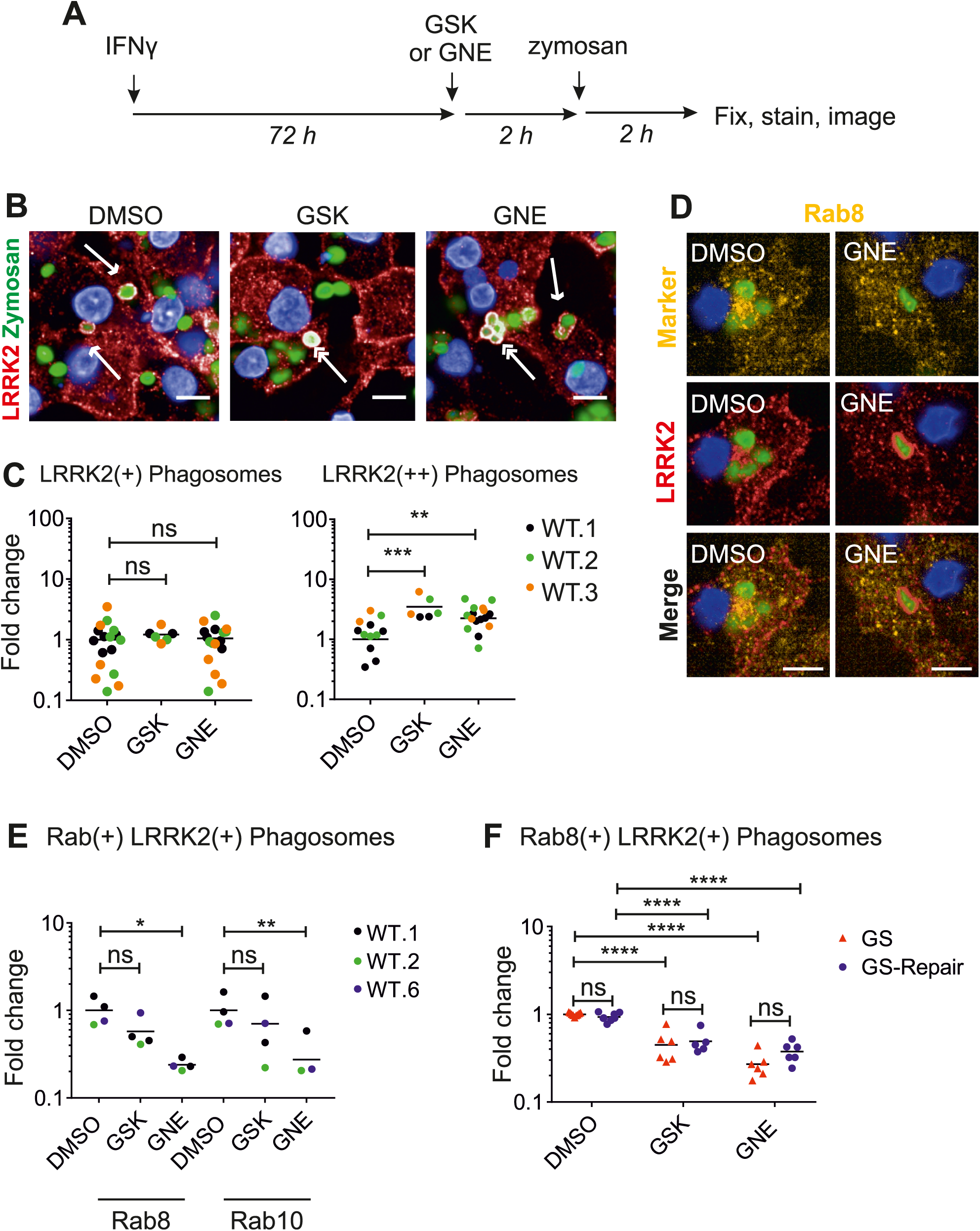
LRRK2 kinase inhibitors increase LRRK2 density at phagosomes, but inhibit Rab8 and Rab10 recruitment. (A) hiPSC-macrophages were pre-treated with GSK (1 µM) or GNE (1 µM) for 2 h before Alexa-488 zymosan bioparticles were added. Phagocytosis proceeded for 2 h then cells were fixed and stained with antibody against LRRK2. (B) Confocal images show examples of LRRK2(+) phagosomes (single headed arrows) and LRRK2(++) (‘supercoated’) phagosomes (double-headed arrows). Scale bars represent 10 µm. (C) Each dot shows data (mean) collected from at least 300 cells based on images acquired from at least five randomized fields. Two-way ANOVA with Fisher’s LSD test was used for the statistical analysis. (D) Confocal images show an example of lack of recruitment of Rab8 to LRRK2(++) phagosome in the presence of LRRK2-kinase inhibitor GNE (right-hand panel), whereas DMSO control shows colocalisation of the two proteins at phagosomes (left-hand panel). (E) Quantification of effect of LRRK2 kinase inhibitors GSK and GNE on recruitment of Rab8 and Rab10 to LRRK2(+) phagosomes. Data from 3 independent hiPSC-derived macrophage lines, each data point represents the mean of 20 fields per independent replicate, 1-way ANOVA, Dunnetts post-hoc comparison, NB fourth data point for Rab10+GNE is below axis. (F) As (E) but data from GS and GS-repair hiPSC-macrophages.

Although LRRK2 kinase inhibition led to an increase in LRRK2 presence at phagosomes, Rab8a and Rab10 colocalisation at LRRK2(+) phagosomes was reduced (Figure 6D,E). This observation was also replicated in the G2019S isogenic pair (Figure 6F). Note that whilst kinase inhibition significantly reduced Rab8a and10 recruitment, there was no significant difference between the G2019S line and its isogenic corrected line, implying again that this monoallelic mutation is subtle in relation to the effects of potent chemical inhibitors. Together, these results show that although kinase inhibition causes LRRK2 to accumulate on phagosomes, without kinase activity LRRK2 cannot recruit Rab8a or Rab10.

## Discussion

Here we have characterised a platform for understanding LRRK2 function using human iPS-derived macrophages and microglia, including gene-edited lines, which has enabled us to interrogate the involvement of LRRK2 in phagocytic processes in authentic, professional phagocytes.

Whilst numerous studies have used hiPSC-derived neurons and astrocytes to study LRRK2 function (reviewed in (Booth, 2017)), the advantages of iPSC technology have barely begun to be applied to studying its function in the myeloid lineage. Our hiPSC-macrophages have been used in one previous study as a validation of observations in mouse that *M.tuberculosis* survival inside macrophages requires LRRK2(Härtlova et al., 2018). To our knowledge, only one other group has previously looked at LRRK2 in hiPSC-derived myeloid cells, describing an impact of the G2019S mutation on differentiation capacity(Speidel et al., 2016) – we did not find this in our system over multiple differentiations with our G2019S isogenic pair, nor with our LRRK2 KO isogenic pair.

We show that endogenous LRRK2 in hiPSC-macrophages and microglia is strongly induced by immune signals, particularly IFNγ, as reported in other systems (Gardet et al., 2010; Lee et al.). This upregulation reveals additional bands by Western blot, notably a prominent band running at ∼170 kD (187kD predicted from proteomic analysis), likely equivalent to the ∼160 kD truncated product reported by others in mouse kidney proximal tubule cells (also detectable in lung and spleen but not brain)(Herzig et al., 2011) and the ∼200 kD product reported in neutrophils and PBMCs (Fan et al., 2018). We have identified the LRRK2 cleavage region in hiPSC-macrophages to be within the ANK-LRR interdomain region, likely more susceptible to cleavage, particularly if 14-3-3 proteins are not bound to the nearby S910/S935 phosphorylation sites. Structural modelling predicts that LRRK2 dimers have C-terminal (kinase and WD40) domains folded backward, in contact with N-terminal domains, suggesting that N-terminal domains may modulate enzymatic activity(Guaitoli et al., 2016). While this cleavage product could be merely the first stage of LRRK2 degradation, it could also, by lacking the N-terminal Armadillo/Ankyrin domains, affect LRRK2 cellular localisation, kinase activity and/or interaction with other proteins.

In our assays, LRRK2 did not impact on the initial uptake of phagocytic particles. Previous studies have yielded conflicting results regarding LRRK2 and phagocytic uptake. siRNA-mediated knock down of LRRK2 has been variously reported to reduce phagocytic uptake in mouse microglial transformed BV-2 cells(Marker et al., 2012) and, conversely, to have no effect in mouse macrophage transformed RAW264.7 cells and BV-2 cells (Schapansky et al., 2014), nor in primary mouse *LRRK2* KO microglia versus WT(Härtlova et al., 2018; Maekawa et al., 2016). A recent study across multiple systems (fly, mouse and Parkinson’s patients) has implicated LRRK2 in phagocytic uptake of beads or *E.coli*, through LRRK2 phosphorylating the actin-remodelling complex component, WAVE2 (Kim et al., 2018). It is not clear why our results diverge from this study, as hiPSC-macrophages express similar levels of WAVE2 to primary human monocytes and microglia(Haenseler et al., 2017a), but it may reflect technical differences (e.g. that study used serum-containing medium, which can lead to engagement of different receptor subsets; and *ex vivo* primary cells may reflect background donor physiological status) that could affect the engagement of the actin-remodelling complex and the relative impact of LRRK2 in the process (Rotty et al., 2017).

Our observations that in hiPSC-macrophages, LRRK2 is recruited to late phagosomes containing zymosan, *E. coli* or *S.typhimurium*, extends the observation made by Gardet et al. (Gardet et al., 2010), who observed recruitment of LRRK2 to *S. typhimurium*-containing phagosomes in RAW264.7 cells. Zymosan engages TLR2 and dectin-1 receptors, *E. coli* engages TLR4, and *S.typhimurium* engages TLR2/TLR4. We did not observe LRRK2 recruitment to α-synuclein fibrils under these experimental conditions. Oligomeric αsyn engages TLR1/2, and monomeric αsyn engages TLR4(Fellner et al., 2013; Kim et al., 2012), but exactly which receptors αsyn fibrils engage is currently unclear. The relationship between LRRK2 and aggregated αsyn is not, therefore, as clear-cut as between LRRK2 and bacterial and fungal pathogens in this phagocytic context.

In our system, LRRK2 colocalises with late phagocytic markers Rab9 and LAMP1, rather than with the early phagocytic marker Rab5, and also colocalises with Rab8a and Rab10, but not with Rab7. Rab9 facilitates the recycling of mannose 6-phosphate receptors (M6PRs) between the phagosomes and the trans-golgi network (TGN). M6PRs are important in delivering newly synthesized lysosomal enzymes from the TGN to phagosomes(Riederer et al., 1994). LAMP-1, while found abundantly on lysosomes(Eskelinen et al., 2003), is recruited to phagosomes prior to Rab7 recruitment, which is required for the formation of phagolysosomes(Huynh et al., 2007). Therefore, it is conceivable that LRRK2 is recruited prior to recruitment of Rab7. Rab8a is involved in vesicle transport and membrane recycling to the cell surface(Banton et al., 2014), autophagy (with the ALS-related gene, C9orf72, implicated as a Rab8 GEF) (Corbier and Sellier, 2017), and has previously been linked to LRRK2 in, variously, endolysosomal trafficking(Rivero-Ríos et al., 2019), centrosome function (Madero-Pérez et al., 2018) and ciliogenesis(Steger et al., 2017). Rab10 has been associated with TLR4 recycling to the surface from endosomes/golgi(Wang et al., 2010), and has also been implicated in LRRK2-mediated ciliogenesis(Steger et al., 2017). Therefore, these Rabs could be recruited by LRRK2 to maturing phagosomes in our system to participate in recycling/rerouting of phagocytosed membrane, receptors and contents. Hartlova et al.,(Härtlova et al., 2018) implicate LRRK2 in retarding *M.tuberculosis* phagosome maturation. Combined with our findings here, it is conceivable that LRRK2 provides links to alternative fates for phagosomal components and contents, other than a direct route to total proteolysis.

Steger et al., identified Rabs as bona fide substrates of LRRK2, with Rab3A/B/C/D, Rab8A/B, Rab10, Rab12, Rab35 and Rab43 as the subset of Rabs phosphorylated by LRRK2 in cells(Steger et al., 2017), and phosphorylation of Rab10 has been exploited as a readout of LRRK2 kinase activity in human neutrophils (Fan et al., 2018). It is likely then, that Rab8a and Rab10, which colocalise with LRRK2 at phagosomes, and whose recruitment is dependent on LRRK2 presence and kinase activity, are being phosphorylated by LRRK2 at this location. During the final phase of this study, Eguchi et al., reported that LRRK2 recruits and phosphorylates these Rabs to chloroquine-induced overload-stressed lysosomes in mouse RAW264.7 cells (briefly also noting LRRK2 recruitment to zymosan phagosomes), leading to release of lysosomal contents (Eguchi et al., 2018). We have extended their observations to show LRRK2-dependent Rab8a and Rab10 recruitment to phagosomes in authentic human macrophages, and have observed LRRK2 ‘supercoating’ of phagosomes upon LRRK2 kinase inhibitor treatment. Accumulation of kinase-inhibited LRRK2 suggests that LRRK2 does not need kinase activity to be recruited to phagosomes, but needs to phosphorylate a substrate in order to leave the phagosome.

Most papers studying LRRK2 function rely on biochemical approaches, Western blot and tagged or overexpressed constructs, which may not localise properly, due to tagging and/or overexpression leading to incorrect stoichiometry with other proteins. By optimising staining for visualising LRRK2 in hiPSC-macrophages by confocal microscopy, we have been able to interrogate the translocation of endogenous, untagged LRRK2 within relevant human cells, revealing the point during phagosome maturation that LRRK2 recruits Rab8a and Rab10, and pointing to a possible role for LRRK2 in rerouting and recycling phagocytosed membrane, receptors and contents.

## Experimental Procedures

### Generation of *LRRK2* modified hiPSC lines

The iPSC lines used in this paper were derived from dermal fibroblasts using non-integrating Sendai reprogramming vectors (Cytotune, Life Technologies), from donors recruited through StemBANCC(Morrison et al., 2015)/Oxford Parkinson’s Disease Centre: participants were recruited to this study having given signed informed consent, which included derivation of hiPSC lines from skin biopsies (Ethics Committee: National Health Service, Health Research Authority, NRES Committee South Central, Berkshire, UK, who specifically approved this part of the study (REC 10/H0505/71). They are deposited in the European Bank for induced pluripotent Stem Cells, EBiSC, https://cells.ebisc.org/ and listed in hPSCreg, https://hpscreg.eu/. See Table S1 for details of lines(Buskin et al., 2018; Dafinca et al., 2016; Fernandes et al., 2016; Haenseler et al., 2017b), and Figure S1 for relevant characterisation data. iPSC were maintained in mTeSR™1 (StemCell Technologies) or E8 (Life Technologies), on hESC-qualified Matrigel-(BD) or Geltrex-(Life Technologies) coated plates, and passaged as clumps with 0.5 mM EDTA in PBS (Beers et al., 2012). Large-scale SNP-QCed frozen batches were used for experiments to ensure consistency.

### Differentiation of macrophages and microglia from hiPSCs

Macrophages and microglia were differentiated using previously published protocols(Haenseler et al., 2017a; van Wilgenburg et al., 2013). See Table S3 for media compositions. In brief, embryoid bodies (EBs) were generated in mTeSR medium with BMP4, VEGF and SCF then differentiated along a primitive myeloid pathway in T175 flasks in X-VIVO 15 medium with M-CSF and IL-3. After about three weeks onwards, macrophage precursors emerging into the supernatant were harvested, passed through a 40 μM cell strainer, centrifuged 5 min at 400 g and further differentiated for 1 week into macrophages in X-VIVO 15 containing 100 ng/mL M-CSF, with a 50% medium change at day 4.

Dual SMAD inhibition was used to differentiate cortical neurons from hiPSC based on Shi et al. (Shi et al., 2012). Day 43 cortical neurons were then cocultured with macrophage precursors in microglia differentiation medium containing IL-34 GM-CSF and used 14 days of differentiation in co-culture as described in Haenseler et al.(Haenseler et al., 2017a).

### Quantification of LRRK2 positive phagosomes

For each experiment, 50,000 macrophage precursors were seeded and differentiated for 1 week in separate wells of Ibidi 96 well µ-plate (Ibidi, 89626). hiPSC-macrophages were treated with IFNγ (100 ng/mL) for 72 h to increase expression of endogenous LRRK2, prior to the addition of the phagocytic materials: Alexa Fluor 488-conjugated zymosan (Life Technologies, Z23373), Alexa Fluor 488-conjugated *Escherichia* coli (Life Technologies, E13231), GFP-expressing *Salmonella typhimurium* (Tocris, NCTC 12023, MM11-25). Oregon Green 488-conjugated αsyn-fibrils (gift from Dr Kelvin Luk, University of Pennsylvania) were prepared with endotoxin-depletion according to the published methods, tested for endotoxin levels and used at a final dilution of ≤0.01 EU/ml endotoxin (a level considered negligible) (Luk et al., 2007; Luk et al., 2009).

After 2 h, cells were washed, fixed and stained as above. Five to 20 Z-stacked confocal images per well were acquired from randomized fields using Opera Phenix High Content Screening System (PerkinElmer) with a 63x objective. Quantification of phagosomes was carried out by Columbus Image Data Storage and Analysis System (CambridgeSoft). Detailed methods used for the analysis is described in supplemental information.

### Flow Cytometry Phagocytosis assays

Uptake of bioparticles was quantitatively assessed by adding Alexa Fluor 488 conjugated zymosan bioparticles (Life Technologies, Z23373) to hiPSC-macrophages (2 bioparticles per cell) for 30 min at 37 °C, followed by wash steps with PBS and trypan blue (250 μg/mL in PBS) to quench non-internalized bioparticles. Cells were detached by TrypLE and gentle manual scraping, centrifuged at 400 *g* for 5 min, and fixed with 4% formaldehyde in PBS. Internalized zymosan particles were quantified using a Becton-Dickinson FACS Calibur flow cytometer (BD biosciences) and analysed using FlowJo software.

### Statistics

Statistical analysis was carried out with GraphPad Prism. All data are represented as mean ± SEM of at least three independent experiments carried out by using cells collected from at least three independent differentiation batches unless stated otherwise. When comparing the means from multiple groups against one control group, one-way analysis of variance (ANOVA) with Dunnett’s *post hoc* comparison, or for data that do not display normal distribution, non-parametrical Kruskal-Wallis with Dunnett’s *post hoc* comparison was used, unless stated otherwise. Statistical significance was defined as ns, *P* > 0.05; \**P* < 0.05; \*\**P* < 0.01; \*\*\**P* < 0.001; \*\*\*\**P* < 0.0001.

## Author Contributions

Conceptualization, H.L., W.S.J., S.A.C.; Methodology, H.L., R.F., P.J.C., F.J.N., M.S., S.A.C.; Investigation, H.L., R.F., I.S., S.A.C.; Formal Analysis, H.L., I.S., S.A.C.; Writing – Original Draft, H.L., S.A.C.; Writing – Review & Editing, H.L., W.S.J., S.A.C.; Funding Acquisition, R.W.-M., W.S.J., S.A.C.; Resources, R.F., P.J.P., M.S., J.V., R.W-M., W.S.J., S.A.C.; Supervision, R.W-M., W.S.J., S.A.C.

## Acknowledgments

Financial support: The Wellcome Trust WTISSF121302 and the Oxford Martin School LC0910-004 (James Martin Stem Cell Facility Oxford, S.A.C.); the MRC Dementias Platform UK Stem Cell Network Capital Equipment MC_EX_MR/N50192X/1, Partnership MR/N013255/1 (R.W-M., S.A.C.) and Momentum MC_PC_16034 (F.J.N., S.A.C.) Awards; ARUK Oxford Network pump-priming award (H.L.). The work was supported by the Innovative Medicines Initiative Joint Undertaking under grant agreement number 115439, resources of which are composed of financial contribution from the European Union’s Seventh Framework Program (FP7/2007e2013) and EFPIA companies’ in kind contribution (R.F.). We thank the High-Throughput Genomics Group at the Wellcome Trust Center for Human Genetics, Oxford (Funded by Wellcome Trust grant reference 090532/Z/09/Z and MRC Hub grant G0900747 91070) for the generation of Illumina genotyping data. We would also like to thank the National Phenotypic Screening Center for instrument support. Samples and associated clinical data were supplied by the Oxford Parkinson’s Disease Center (OPDC) study, funded by the Monument Trust Discovery Award from Parkinson’s UK, a charity registered in England and Wales (2581970) and in Scotland (SC037554), with the support of the National Institute for Health Research (NIHR) Oxford Biomedical Research Center based at Oxford University Hospitals NHS Trust and University of Oxford, and the NIHR Comprehensive Local Research Network (J.V., R.W-M., S.A.C.).

