## Supplemental Information for "LRRK2 is recruited to phagosomes and co-recruits Rab8 and Rab10 in human pluripotent stem cell-derived macrophages"

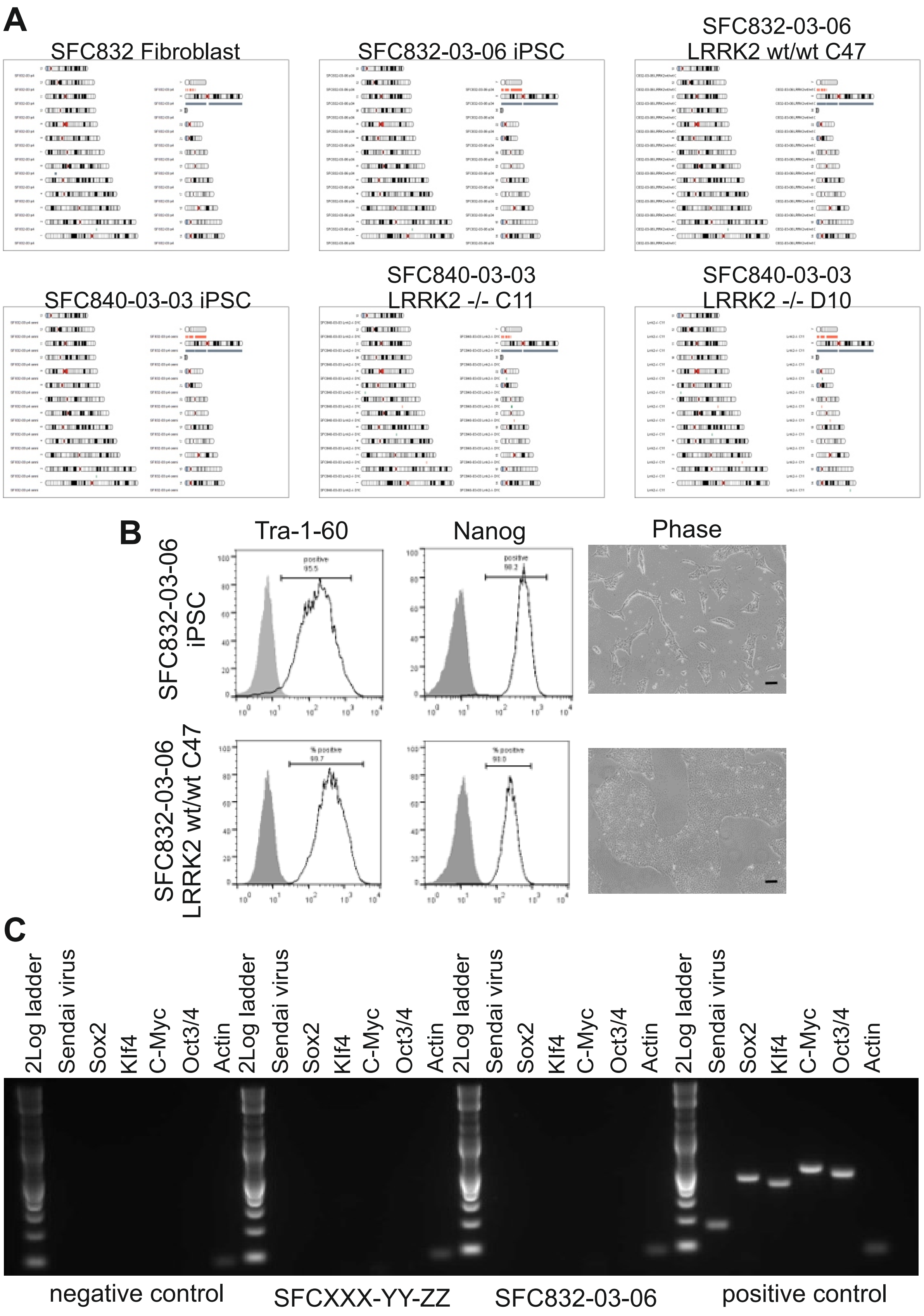
A
SFC832 Fibroblast
SFC832-03-06 iPSC
SFC832-03-06
LRRK2 wt/wt C47
SFC840-03-03 iPSC
SFC840-03-03
LRRK2 -/- C11
SFC840-03-03
LRRK2 -/- D10
B
Tra-1-60
Nanog
Phase
SFC832-03-06
iPSC
SFC832-03-06
LRRK2 wt/wt C47
C
2Log ladder
Sendai virus
Sox2
Klf4
C-Myc
Oct3/4
Actin
negative control
SFCXXX-YY-ZZ
SFC832-03-06
positive control

**Figure S1. Quality control of hiPSC lines used in this study (related to Figure 1 and to Experimental procedures, Generation of LRRK2 modified hiPSC lines).**

HiPSC line SFC832-03-06 and gene-edited lines characterised in this manuscript. (A) Genome integrity of SFC832-03-06 and gene-edited lines, assessed by Illumina Human CytoSNP12v2.1 or OmniExpress24 SNP array. Karyograms (KaryoStudio, Illumina) show amplifications (green)/deletions (orange)/LOH regions (grey) alongside the relevant chromosome (NB two X chromosomes annotated grey, Y chromosome orange, with poor coverage by CytoSNP12v2.1 versus OmniExpress24). (B) FACS analysis confirms expression of pluripotency markers Tra-1-60 and Nanog; open black plot represents antibody, filled grey plot is isotype control; Right-hand panel shows expected hiPSC colony morphology (phase-contrast microscopy), Scale bar, 100 µm. (C) Clearance of Cytotune1 Sendai vectors from SFC832-03-06 (Log2 ladder; Sendai backbone 181bp; Sox2 451 bp; Klf4 410 bp; c-myc 532 bp; Oct-4 483 bp; β-actin control 92 bp; + control is fibroblasts infected with Cytotune 5 d previously).

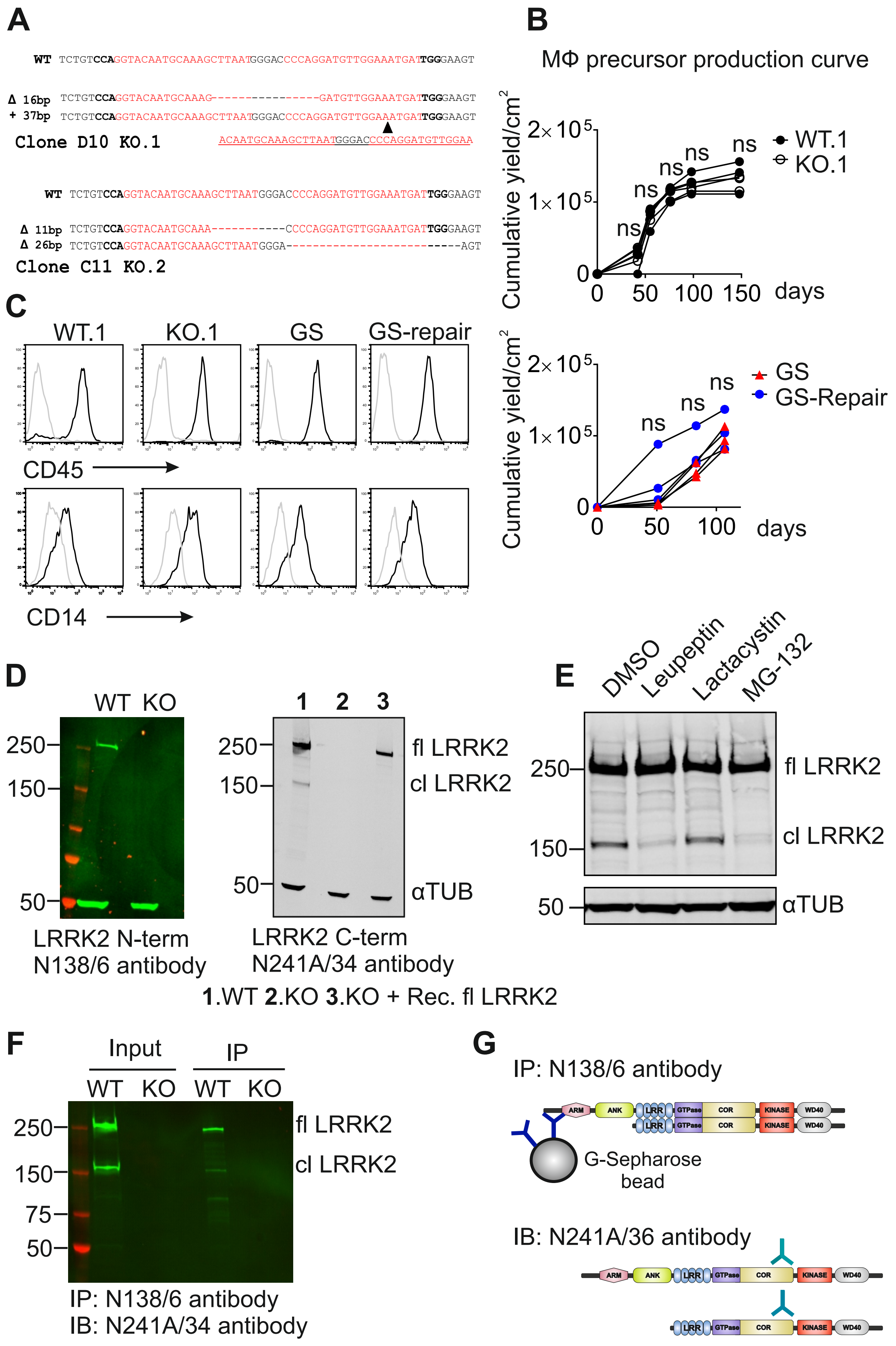
A
WT TCTGTCCAGGTACAATGCAAAGCTTAATGGGACCCCAGGATGTTGGAAATGATTGGGAAGT
Δ 16bp TCTGTCCAGGTACAATGCAAAG----------------GATGTTGGAAATGATTGGGAAGT
+ 37bp TCTGTCCAGGTACAATGCAAAGCTTAATGGGACCCCAGGATGTTGGAAATGATTGGGAAGT
Clone D10 KO.1
ACAATGCAAAGCTTAATGGGACCCCAGGATGTTGGAA
WT TCTGTCCAGGTACAATGCAAAGCTTAATGGGACCCCAGGATGTTGGAAATGATTGGGAAGT
Δ 11bp TCTGTCCAGGTACAATGCAAA-----------CCCCAGGATGTTGGAAATGATTGGGAAGT
Δ 26bp TCTGTCCAGGTACAATGCAAAGCTTAATGGGA--------------------------AGT
Clone C11 KO.2
B
MΦ precursor production curve
Cumulative yield/cm²
2×10⁵
1×10⁵
0
0 50 100 150 days
ns
WT.1
KO.1
GS
GS-Repair
0 50 100 days
C
WT.1
KO.1
GS
GS-repair
CD45
CD14
D
WT KO
250
150
50
LRRK2 N-term
N138/6 antibody
1 2 3
fl LRRK2
cl LRRK2
αTUB
LRRK2 C-term
N241A/34 antibody
1.WT 2.KO 3.KO + Rec. fl LRRK2
E
DMSO
Leupeptin
Lactacystin
MG-132
250
150
50
fl LRRK2
cl LRRK2
αTUB
F
Input
IP
WT KO WT KO
250
150
75
50
fl LRRK2
cl LRRK2
IP: N138/6 antibody
IB: N241A/34 antibody
G
IP: N138/6 antibody
ARM
ANK
LRR
GTPase
COR
KINASE
WD40
G-Sepharose bead
IB: N241A/36 antibody

**Figure S2. Differentiation of LRRK2 gene-edited iPSC to macrophages and analysis of LRRK2 species (related to Figure 1).**

(A) Sequences of each allele of hiPSC clones with CRISPR/Cas9-mediated double-allelic knockout of LRRK2. Black triangle indicates position of an inserted sequence. (B) Cumulative yield of macrophage precursors harvested from the supernatant of iPSC-macrophage differentiation cultures, for final plating for mature macrophages. Each line represents an independent differentiation. Two-way ANOVA with Bonferroni multiple comparisons test was used for statistical analysis. (C) Flow cytometry of macrophage precursor cells from LRRK2 gene-edited hiPSC lines, for key myeloid markers CD45 and CD14. (D) Left panel, western blot of hiPSC-macrophage whole-cell lysate, probed with the N-terminal antibody for LRRK2, showing lack of the 150kD band. Right panel, as left panel, but probed with C-terminal antibody for LRRK2 and with final column of LRRK2 KO macrophage lysate spiked with recombinant LRRK2, demonstrating lack of degradation of the recombinant protein in the experimental cell lysis conditions used in this study. (E) Western blot of whole cell lysates of hiPSC-macrophages incubated with protease inhibitors, probed with C-terminal LRRK2 antibody N241A/34. Lower panel, quantification of above, mean and SEM of 3 experiments, 1-way ANOVA. (F) Western blot of immunoprecipitation of LRRK2 from hiPSC-macrophages using N-terminal antibody, and probed with C-terminal antibody, showing co-precipitation of C-terminal truncated product. D-F, all with 72 h IFNγ activation.

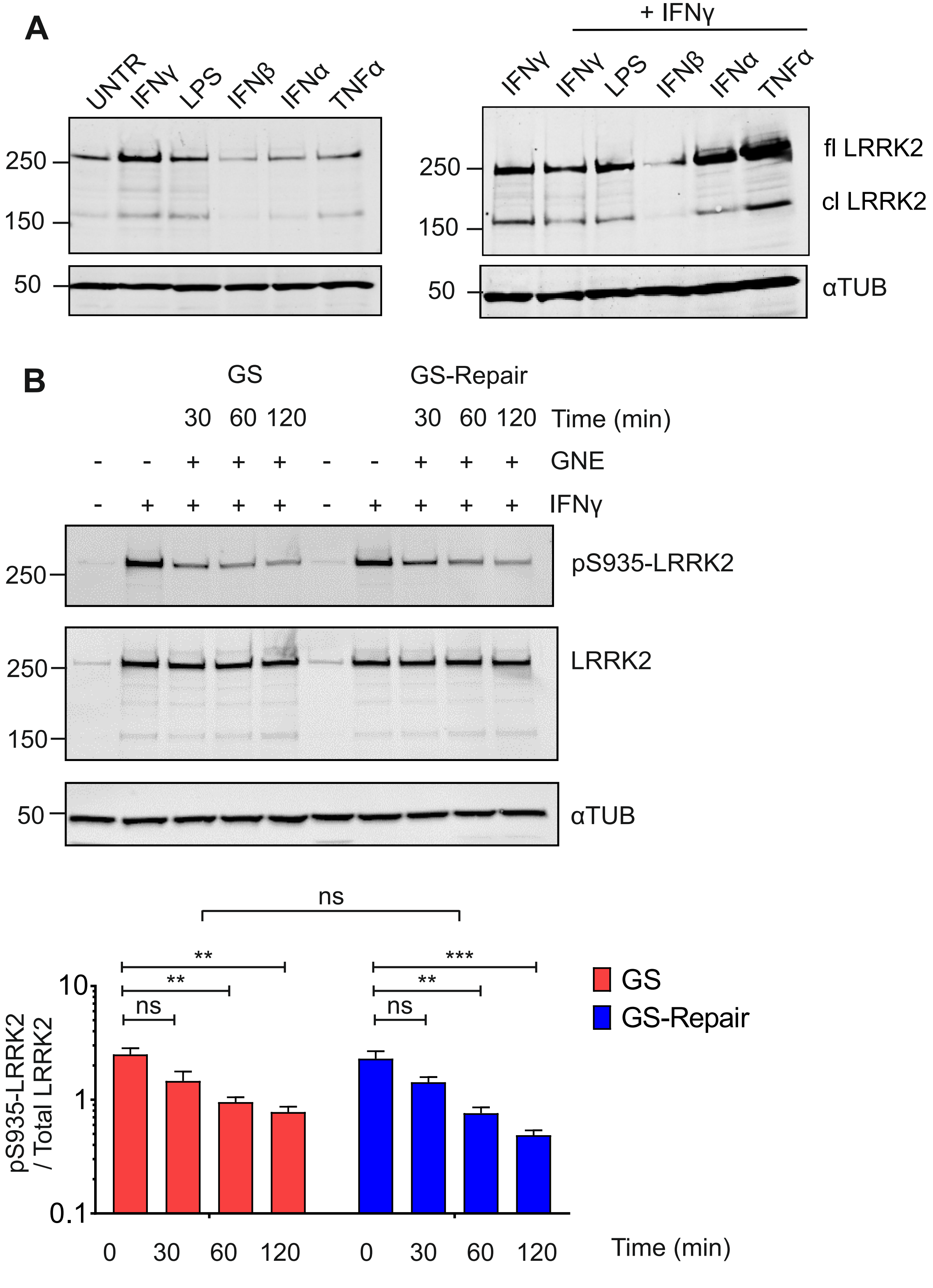
A
UNTR
IFNγ
LPS
IFNβ
IFNα
TNFα
250
150
50
+ IFNγ
fl LRRK2
cl LRRK2
αTUB
B
GS
GS-Repair
30 60 120
Time (min)
GNE
IFNγ
pS935-LRRK2
LRRK2
ns
**
***
GS
GS-Repair
pS935-LRRK2 / Total LRRK2
10
1
0.1
0
30
60
120

**Figure S3. LRRK2 protein expression in activated hiPSC-macrophages, and with LRRK2 kinase inhibition (related to Figures 2 and 3).**

(A) Western blot of hiPSC-macrophage whole-cell lysates following 72 h activation as indicated, probed with C-terminal LRRK2 antibody N241A/34. (B) As (A), for the LRRK2 G2019S and corrected isogenic cell line pair, with activation with IFNγ and incubation with LRRK2 kinase inhibitor GNE for 2 hours prior to cell lysis, plus additional WB probing with antibody for phospho-S935 LRRK2. Lower panel, quantification of above, with mean and SEM of 3 experiments. Statistical analysis, 1-way ANOVA

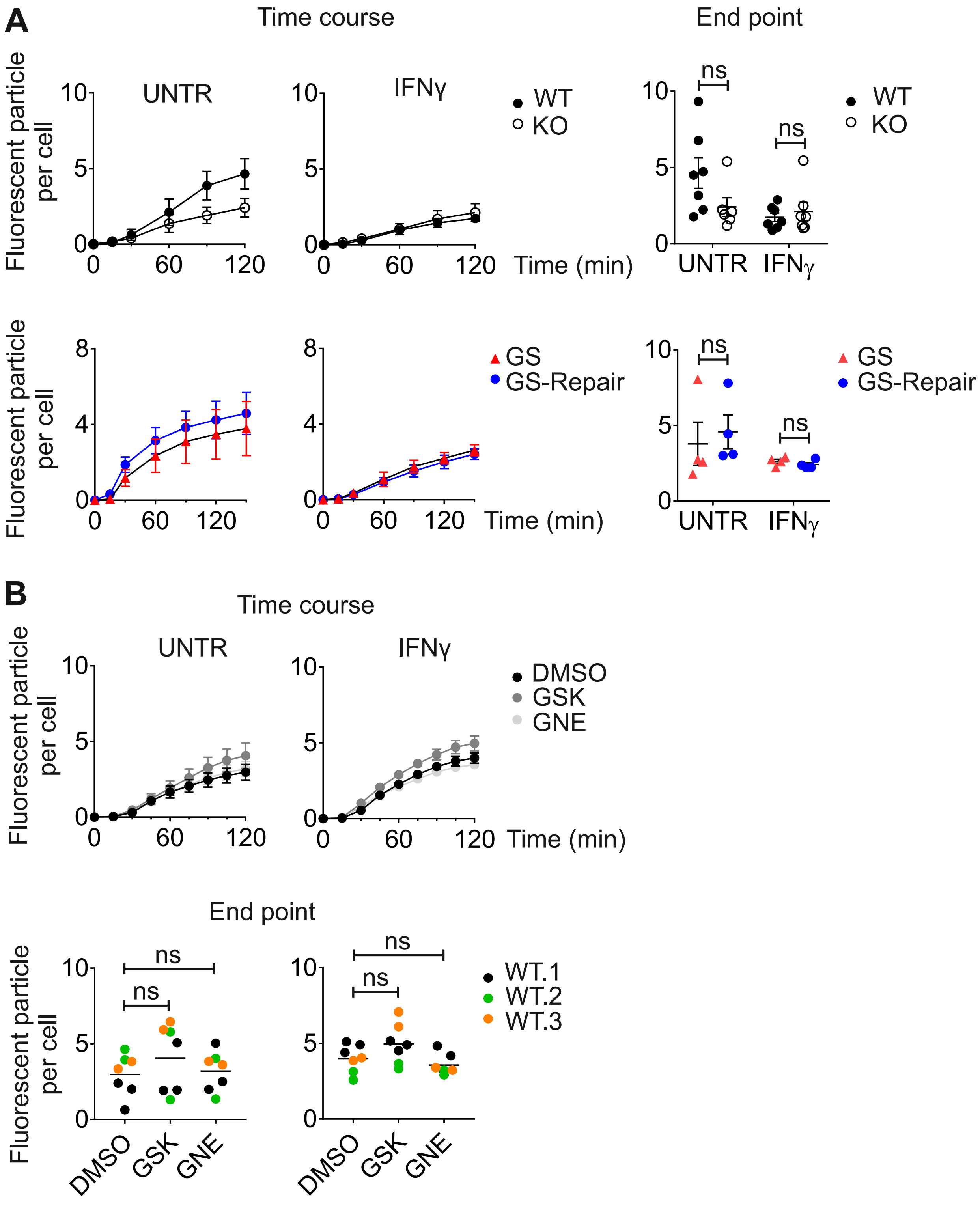


**Figure S4. Acidification of phagosomes in hiPSC-macrophages (related to Figure 3).**

(A) Time-course of uptake of pHrodo-labelled zymosan by LRRK2 gene-edited hiPSC-macrophages, quantified by IncuCyte ZOOM Software (Essenbioscience). Right-hand panel, statistical analysis, mean and SEM., Student t-test. (B) As (A) but with WT hiPSC-macrophages treated with LRRK2 kinase inhibitors GSK or GNE for 2 h before adding pHrodo-zymosan.

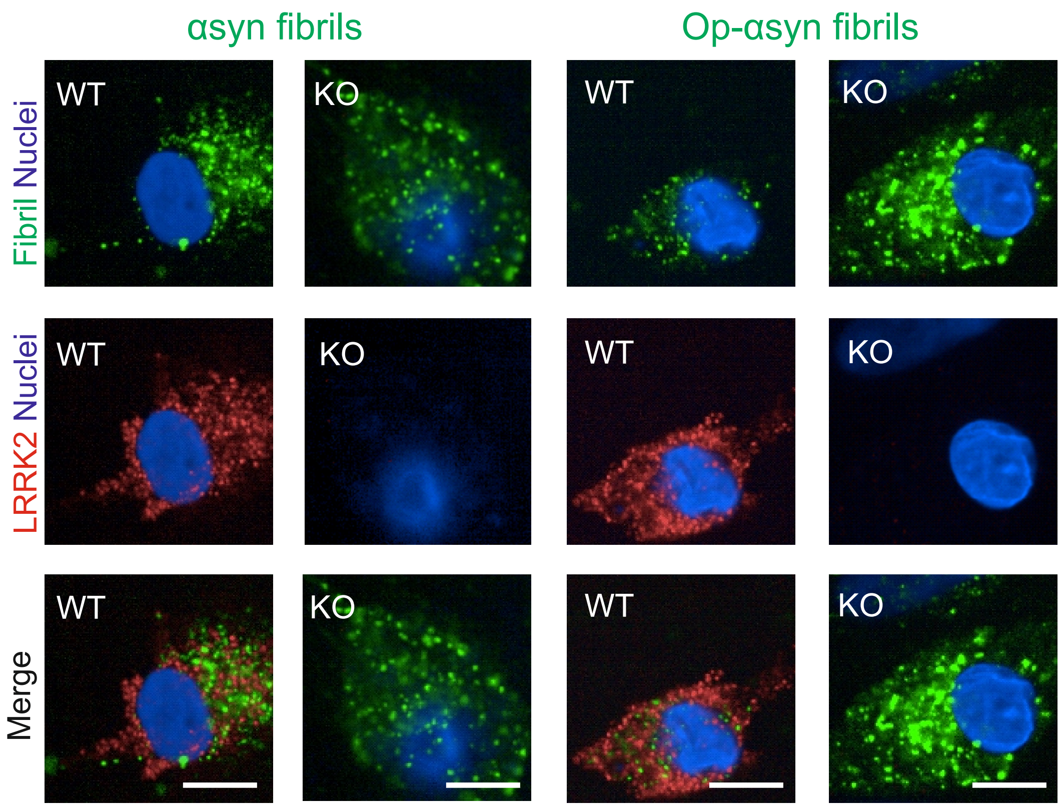


**Figure S5. Additional phagocytic meal LRRK2-localisation data (related to Figure 4).**

Confocal images of hiPSC-macrophages following uptake of α-synuclein fibrils (Alexa-488 labelled, with and without opsonisation with pooled human serum), stained for LRRK2 (Alexa-647, red). Scale bar, 10 µm.

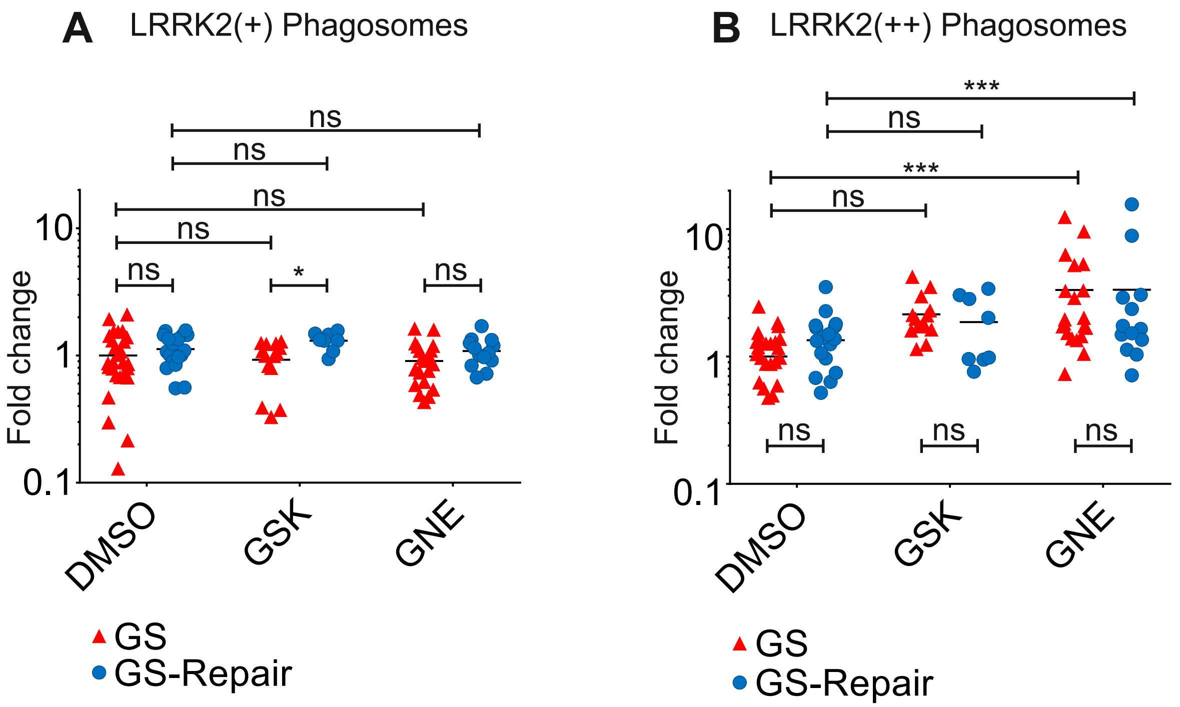


**Figure S6. LRRK2 recruitment to hiPSC-macrophage phagosomes with LRRK2 kinase inhibition in LRRK2 G2019S isogenic pair (related to Figure 6).**

(A) Quantification of number of LRRK2(+) phagosomes in LRRK2 G2019S isogenic pair, as in Figure 6F. (B) As (A) but LRRK2(++) ('supercoated') phagosomes. Each data point represents the mean of 20 fields per independent replicate, 1-way ANOVA, Dunnett's post-hoc comparison.

**Table S1. Human iPSC lines.**

| Labels | | ID of fibroblast | ID of iPSC clone | Diagnosis | Genotype LRRK2 | Gender | Age of Biopsy (years) | Repro-gramming method | PMID |
|---|---|---|---|---|---|---|---|---|---|
| WT.1 | ● | SFC840 | SFC840-03-03 | Healthy | WT/WT | F | 67 | Cytotune1 | 26905200 |
| WT.2 | ● | SFC856 | SFC856-03-04 | Healthy | WT/WT | M | 78 | Cytotune2 | 28827786 |
| WT.3 | ● | SBAD2 | SBAD2-01 | Healthy | WT/WT | M | 51 | Cytotune1 | 30315276 |
| WT.4 | ● | SFC180 | SFC180-01-01 | Healthy | WT/WT | F | 60 | Cytotune2 | 28827786 |
| WT.5 | ● | SFC841 | SFC841-03-01 | Healthy | WT/WT | M | 36 | Cytotune1 | 27097283 |
| WT.6 | ● | SFC854 | SFC854-03-02 | Healthy | WT/WT | M | 72 | Cytotune2 | 28827786 |
| KO.1 | ○ | SFC840 | SFC840-03-03 $LRRK2^{-/-}$ D10 | CRISPR/ Cas-9 edited | -/- | | | | 29789389 uses this clone; This paper for characterisation data |
| KO.2 | ○ | SFC840 | SFC840-03-03 $LRRK2^{-/-}$ C11 | CRISPR/ Cas-9 edited | -/- | | | | This paper |
| GS | ▲ | SFC832 | SFC832-03-06 | PD | G2019S/WT | F | 77 | Cytotune1 | This paper |
| GS-Repair | ● | SFC832 | SFC832-03-06 $LRRK2^{WT/WT}$ C47 | CRISPR/ Cas-9 edited | WT/WT | | | | This paper |

**Table S2. Guide RNA sequences.**

| | **Sequences 5' – 3'** |
|---|---|
| LRRK2 KO gRNA oligo1 | CACCGATTAAGCTTTGCATTGTACC |
| LRRK2 KO gRNA oligo2 | AAACGGTACAATGCAAAGCTTAATC |
| LRRK2 KO gRNA oligo1 | CACCGCCCAGGATGTTGGAAATGAT |
| LRRK2 KO gRNA oligo2 | AAACATCATTTCCAACATCCTGGGC |
| LRRK2 G2019S Repair gRNA oligo 1 | CACCGTCAGCAATCTTTGCAATGA |
| LRRK2 G2019S Repair gRNA oligo 2 | AAACTCATTCGAAAGATTGCTGAC |
| LRRK2 G2019S Repair gRNA oligo 1 | CACCGTCAGTACTGCTGTAGAATG |
| LRRK2 G2019S Repair gRNA oligo 2 | AAACCATTCTACAGCAGTACTGAC |

**Table S3. Differentiation media.**

| Media | Composition |
|---|---|
| **Macrophage differentiation** | |
| EB differentiation medium | mTeSR (StemCell Technologies, 12491), 50 ng/mL BMP4 (Gibco, PHC9534), 50 ng/mL VEGF (PeproTech, 100-20), 20 ng/mL SCF (Miltenyi Biotec, 130-094-303) |
| Factory medium | X-VIVO-15 (Lonza, BE04-418F), 100 ng/mL M-CSF (Gibco, PHC9501), 25 ng/mL IL-3 (Gibco, PHC0033), 2 mM Glutamax, (Gibco 35050035) 100U/mL P/S, 0.055 mM 2-ME |
| Macrophage differentiation medium | X-VIVO-15, 100 ng/mL M-CSF, 2 mM Glutamax, 100 U/mL P/S, 0.055 mM 2-ME |
| **Microglia-cortical neuron co-culture** | |
| Neural induction medium | Neural maintenance medium, 10 µM SB431542 (Tocris, 1614), 1 µM Dorsomorphin (Tocris, 3093) |
| Neural maintenance medium | 50:50 mix: DMEM F12 (Gibco, 21331020) supplemented 1 in 100 N2 (Gibco, 17502048), 2 mM Glutamax and Neurobasal medium (Gibco, 21103049) supplemented 1 in 50 B27 (Gibco, 17504044), 2 mM Glutamax |
| Microglia-neuron co-culture medium | Advanced DMEM F12 (Gibco, 12634010), 1 in 100 N2, 100 ng/mL IL-34, (PeproTech, 200-34), 10 ng/mL GM-CSF, 2 mM Glutamax |

**Table S4. Antibodies.**

| | Supplier | Cat No. | WB | IF |
|---|---|---|---|---|
| IBA1 | Abcam | ab5076 | | 1:200 |
| LAMP-1 | Cell Signaling Technology | D2D11 | | 1:200 |
| LRRK2 N-Term (N138/6) | NeuroMAB | 75-188 | 1:1000 | |
| LRRK2 C-Term (N241A/34) | NeuroMAB | 75-253 | 1:1000 | 1:1000 |
| LRRK2 phospho S935 | Abcam | ab133450 | 1:500 | |
| MAP2 | Abcam | ab32454 | | 1:1000 |
| pan-14-3-3 | Thermo Fisher | 51-0700 | | 1:100 |
| Rab7 | Cell Signaling Technology | 9367 | | 1:200 |
| Rab8a | Cell Signaling Technology | D22D8 | | 1:200 |
| Rab9 | Cell Signaling Technology | 5118 | | 1:200 |
| Rab10 | Cell Signaling Technology | D36C4 | | 1:200 |
| α-Tubulin | Sigma | T5168 | 1:5000 | |
| IRDye 680RD Goat anti-Rabbit IgG | Li-Cor | 926-68071 | 1:10,000 | |
| IRDye 800CW Goat anti-Mouse IgG | Li-Cor | 926-32210 | 1:10,000 | |
| Donkey anti-mouse Alexa 647 | Life Technologies | A31571 | | 1:500 |
| Donkey anti-rabbit Alexa 568 | Life Technologies | A10042 | | 1:500 |

### Supplemental Experimental Procedures

#### Gene editing of hiPSC

A double nickase strategy was employed to generate homozygous *LRRK2* KO hiPSC lines. First, gRNAs (Table S2) were cloned into Cas9 nickase plasmid with puromycin section (PX462; Addgene, #48141). hiPSCs were transfected by neon-transfection system (Life Technologies) and were seeded in single cell onto 10 $cm^2$ tissue-culture treated dish (Fisher) at low density to ensure selection of each individual colony. To screen possible KO clones, genomic DNA was extracted and concentration measured using picogreen (Life Technologies). PCR amplicons were produced by Phusion Hot start DNA polymerase (NEB) on PT-200 PCR machine (MJ Research) and quantitative reverse transcript polymerase chain reaction (qRT-PCR) was carried out with LC Green Plus+ (BioChem) and AmpliTaq master mix (Applied Biosystems) on StepOne plus (Applied Biosystems). Continuous melting step was allowed to screen for aberrant melt curve by using High Resolution Melting Software (Applied Biosystems), and the presence of out-of-frame deletion/insertion was confirmed by sequencing analysis. Similarly, for the repair of heterozygous *LRRK2* G2019S mutation, a donor template was designed with silent mutations in the PAM site to maximize the efficiency of gene-editing and *PstI* site for screening clones.

#### Western blot

Cells were washed with ice-cold PBS and then lysed directly by adding lysis buffer (50 mM Tris-HCl, 150 mM NaCl, 0.5 mM EDTA, 1 tablet of protease inhibitor (Roche), 100 µL of phosphatase inhibitor, 1 % maltoside) to a final concentration of $1x10^5$ cells/µL. Cell lysates were centrifuged at 17,000 rpm for 20 min at 4°C. Pellets were discarded and supernatants were mixed with 4x NuPAGE LDS (lithium dodecyl sulphate) sample buffer and 10x NuPAGE sample reducing agent and were heated for 10 min at 70°C. Each sample containing 25 – 30 µg of protein was loaded into either 3-8% Tris-Acetate or 4-12% Bis-Tris NuPAGE pre-cast gels. Electrophoresis was performed for 60 min at 120 V using NuPAGE SDS running buffer. Proteins were transferred onto polyvinylidene difluoride (PVDF) membrane using Pierce Power blotter system (Thermo Scientific). iBind Flex system (Life Technologies) was used to blot membranes with antibodies, using iBind Fluorescent Detection (FD) solution kit according to the manufacturer's instructions. The membrane was scanned with Odyssey Sa Infrared Imaging System (Li-Cor) and fluorescence intensity quantified using MyImage analysis software (Thermo Scientific). See Table S3 for antibodies used.

#### Immunoprecipitation

For crosslinking antibodies to beads, Protein G Sepharose beads (Sigma, P3296) were washed with PBS and incubated with antibody against LRRK2 (NeuroMAB; N241A/34 or N138/6) overnight at 4°C on a rotating wheel. For each preparation of beads, 40 µg of antibodies and 60 µL of Protein G Sepharose beads were mixed in 1 mL PBS. Any unbound antibodies were washed off with 1 mL of 0.2 M sodium borate buffer (pH 9.0). Crosslinking was carried out by incubating antibody-bead complexes with 20 mM dimethyl pimelimidate dihydrochloride (DMP; Sigma, D8388) in 0.2 M sodium borate buffer (pH 9.0) for 40 min at RT. Antibody-bead complexes were washed once with 0.2 M ethanolamine (pH 8.0) and were incubated in 0.2 M ethanolamine buffer for 2 h at RT. Beads were quenched with 0.58% v/v acetic acid and 150 mM NaCl. Crosslinked beads were washed three times with PBS and stored in 4°C until used.

Nonspecific proteins were first cleared by incubating whole cell lysates with naked G Sepharose beads on a rotating wheel in 4°C for 30 min. About 20 x $10^6$ to 30 x $10^6$ cells were incubated in 30 µL of G sepharose beads. Pre-cleared cell lysates were then incubated with G Sepharose beads cross-linked with LRRK2 antibody (NeuroMAB) overnight on a rotating wheel at 4°C. Unbound proteins were washed off three times using lysis buffer without detergents. Finally, LRRK2 interactors were eluted by boiling protein-bead complexes at 70°C for 10 min in SDS elution buffer. See Supplemental information for Mass Spectometry methodology.

#### Immunostaining

Cells were fixed with 4% Paraformaldehyde in PBS in RT for 10 min and were blocked and permeabilized overnight at 4 °C in blocking buffer containing 5% BSA (Sigma, A7906) and 10% normal donkey serum (Sigma, D9663) and 0.1% Triton-X in PBS. Primary antibody was diluted in blocking buffer and cells were stained for 1 h in RT. See Table S3 for antibodies used.

Unbound antibodies were washed in PBS with 0.3% Triton-X three times, 15 min each. Secondary antibodies (Life Technologies) were diluted 1:500 in blocking buffer containing 0.05% Triton-X and incubated for 1 h at RT. Unbound antibodies were again washed in PBS with 0.3% Triton-X for 15 min, three times. Finally, cells were stained with DAPI in PBS with 0.3% Triton-X for 10 min, and washed three times with PBS. Confocal images were taken by Olympus Fluoview FV1200 (Olympus) or an Opera Phenix High-Content Screening System (PerkinElmer).

**Quantification of LRRK2(+) phagosomes using Columbus Image Data Storage and Analysis System**

Analysis of confocal images acquired by OperaPhenix (PerkinElmer) began with “Input image” command, which processed z-stack images by stacking with maximum projection. “Find Nuclei” Method B identified the population of nuclei in DAPI channel. “Find Surrounding Region” using Alexa 647 channel defined approximate cell boundaries, ensuring that only internalised zymosan bioparticles are quantified. “Find Spots” identified zymosan bioparticles using Alexa 488 Channel, with a size range defined. “Find Surrounding Region” found regional spots surrounding each identified zymosan bioparticle. “Calculate Intensity Properties” command calculated intensity of LRRK2 signal (Alexa 647 channel) surrounding zymosan bioparticles. “Select Population” quantified the number of LRRK2(+) phagosomes and also the number of LRRK2 super-coated phagosomes by setting thresholds of signal intensity. The same workflow was used to quantify the number of LRRK2(+) phagosomes that displayed positivity for Rab5, 8a, 9, 10 and LAMP-1 (collectively referred to as markers). But in this case, “Calculate Intensity Properties” command was used twice, first to measure the intensity of LRRK2 signal (Alexa 647) and then for the intensity of the marker signal (Alexa 568) within the LRRK2(+) population (or in whole population for the LRRK2-/- experiments).

**Real-time phagocytosis assay using pHrodo-labelled zymosan bioparticles**

Quantification of acidified phagosomes was evaluated using pHrodo Green zymosan bioparticles (Thermo, P35365). Bioparticles were resuspended in live imaging solution (Thermo, A14291DJ) according to manufacturer's instruction and were sonicated for 10 min (Bioruptor) prior to each experiment. During the assay, 70,000 hiPSC-macrophages in each well of a 96-well clear bottom black plate (Corning, CLS3603) were kept in live imaging solution along with 0.2 mg/mL pHrodo bioparticles. The number of fluorescent bioparticles were monitored every 10 min in InCucyte Live Cell Analysis System (Essen Bioscience) for 2 h. To account for any cell number variability, cells were stained NUCLEAR-ID® Red DNA stain (Enzo Life Sciences) at the end of the phagocytosis experiment. For each acquired image, the total number of fluorescent particles was normalized to the total cell number.

**Mass Spectrometry**

Samples were trypsin digested and desalted using X18 tips. Samples were analysed on an Ultimate 3000 RSLCnano HPLC (Dionex) system run in direct injection mode coupled to a QExactive Orbitrap mass spectrometer (Thermo Electron). Samples were resolved on a 25cm x 75μm inner diameter picotip analytical column (New Objective) which was packed in-house with ProntoSIL 120-3 C18 Ace-EPS phase, 3μ bead (Bischoff Chromatography). The system was operated at a flow-rate of 300nL $min^{-1}$. A 120 min gradient was used to separate the peptides. The mass spectrometer was operated in a “Top 10” data dependent acquisition mode. Precursor scans were performed in the orbitrap at a resolving power of 70,000, from which the ten most intense precursor ions were selected by the quadrupole and fragmented by HCD at normalised collision energy of 28%. The quadrupole isolation window was set at *3 m/z*. Charge state +1 ions and undetermined charge state ions were rejected from selection for fragmentation. Dynamic exclusion was enabled for 40s. Data were converted from .RAW to .MGF using ProteoWizard (Chambers et al., 2012). Data was collected and analysed on the Central Proteomics Facilities Pipeline (CPFP)(Trudgian et al., 2010).
